# Rhythmic neural spiking and attentional sampling arising from cortical receptive field interactions

**DOI:** 10.1101/252130

**Authors:** Ricardo Kienitz, Joscha T. Schmiedt, Katharine A. Shapcott, Kleopatra Kouroupaki, Richard C. Saunders, Michael C. Schmid

**Affiliations:** Ernst Strüngmann Institute (ESI) for Neuroscience in Cooperation with Max Planck Society, Deutschordenstraße 46, 60528 Frankfurt a. M., Germany; Epilepsy Center Frankfurt Rhine-Main, Center of Neurology and Neurosurgery, Goethe University, Schleusenweg 2-16, 60528 Frankfurt a.M., Germany; Laboratory of Neuropsychology, NIMH, Convent Drive 49, Bethesda, MD 20892, USA; Institute of Neuroscience, Newcastle University, Framlington Place, Newcastle upon Tyne, NE2 4HH, UK; lead contact

## Abstract

Growing evidence suggests that distributed spatial attention may invoke theta (3-9 Hz) rhythmic sampling processes. The neuronal basis of such attentional sampling is however not fully understood. Here we show using array recordings in visual cortical area V4 of two awake macaques that presenting separate visual stimuli to the excitatory center and suppressive surround of neuronal receptive fields elicits rhythmic multi-unit activity (MUA) at 3-6 Hz. This neuronal rhythm did not depend on small fixational eye movements. In the context of a distributed spatial attention task, during which the monkeys detected a spatially and temporally uncertain target, reaction times (RT) exhibited similar rhythmic fluctuations. RTs were fast or slow depending on the target occurrence during high or low MUA, resulting in rhythmic MUA-RT cross-correlations at at theta frequencies. These findings suggest that theta-rhythmic neuronal activity arises from competitive receptive field interactions and that this rhythm may subserve attentional sampling.

**Highlights:**

- Center-surround interactions induce theta-rhythmic MUA of visual cortex neurons
- The MUA rhythm does not depend on small fixational eye movements
- Reaction time fluctuations lock to the neuronal rhythm under distributed attention

## Introduction

Spatial attention can exhibit fluctuations that under some tested conditions might be rhythmic. In vision, this is apparent for example during overt saccadic exploration of visual scenes during which periods of fixation tend to occur every ∼200 ms, i.e. in the slow theta range [1]. Similar rhythmic exploration is sometimes also observed during apparent fixation periods when subjects perform fast fixational eye movements (microsaccades) [1,2]. Such rhythmic sampling phenomena appear to be not limited to overt behavior, but have also been discovered in investigations on covert distributed spatial attention, i.e. in the absence of overt eye movements. Here the subject’s capacity to detect a change in one of multiple objects is assessed as a function of trial-by-trial varying target onset times. A convergent finding across several recent studies is a theta (3-9 Hz) rhythmic sampling that can be observed in performance or reaction time measures during such distributed attention conditions [3–5]. For example, in a study by Fiebelkorn et al. [6] subjects had to detect a target on one of three possible target positions whereby two positions belonged to the same underlying object and one position to an alternative object. The results confirmed a theta-rhythmic modulation of detection performance under these task conditions. The phase of the rhythm depended on the target location, such that faster reaction times to one location alternated with those to the alternative location of the same object. It therefore appears that the brain might engage a spatial sampling mechanism that operates in the theta range when two or more objects are simultaneously monitored.

EEG/MEG studies in humans confirmed the presence of rhythmic oscillatory responses during a variety attentional tasks [4,7–11]. For example, theta oscillations were measured over visual cortex during tasks that require the tracking of multiple objects, where theta appeared to influence the detectability of a visual target (e.g. [4,9,11,12]). Reports from intracranial neuronal recordings in fixating monkeys sometimes contain theta rhythmic activation in V4 and inferotemporal cortex (IT) [13–18], but the mechanism generating this rhythm and its possible relationship to attentional sampling remain unclear. Two recent studies linked theta and gamma oscillations to microsaccade-occurrences [19,20]. However, it is unclear whether microsaccades constitute a prerequisite for the neuronal rhythm to emerge or whether it can also occur independently in which case the neuronal mechanism generating the rhythm still remains unknown. To address this, we recorded multi-unit activity from V4 neurons while monkeys performed a task invoking attentional sampling. We focused on area V4 as neuronal activity of this area is known to be well associated with attention: Lesions of V4 result in an attentional stimulus selection deficit [21]. The firing of many V4 neurons is modulated by (micro-)saccades [20,22,23] and increased when attention is covertly focused on a stimulus in their receptive field (RF) [24,25]. When in addition to a stimulus in the RF, a second stimulus is added to the RF surround, the neuron’s response is usually suppressed relative to its response to the center stimulus alone [26–29]. Focusing attention to the RF center or surround stimulus will enhance or diminish this surround suppression respectively [25,29–32]. We reasoned that during longer stimulus presentation times the spatial structure of V4 neuron RFs into excitatory center and inhibitory surround might provide the balance of excitation and inhibition that is required for the emergence of oscillatory activity [33–35], which in turn could constitute the neuronal basis for attentional sampling [12]. To investigate this at the level of neighboring neuronal populations we implanted “Utah” micro-electrode arrays into V4 and measured multi-unit activity (MUA, [36]) from the array’s 64 electrodes (see Methods). We first investigated how center-surround RF stimulation could evoke theta rhythmic MUA. As microsaccades, sometimes occurring every 250-300 ms, have been linked with rhythmic neural responses [19,20,37] we also tested their potential contribution to the emergence of theta oscillations. In a final step we investigated the relationship between theta rhythms in the MUA to the monkeys’ reaction times during an attentional detection task.

## Results

### RF center and surround interactions induce theta-rhythmic MUA

To quantify surround suppression in V4 during the passive fixation task, we determined the extent and strength of surround suppression in V4 by systematically increasing the diameter of a disk stimulus presented to a given RF [28]. Stimuli were displayed for 1 second to provide sufficient time to detect oscillatory activity. In all electrodes (40/40 and 57/57 in monkey K and H, respectively) increasing disk size resulted in a response increase up to a maximal response at 1-2° visual degrees (defined as the excitatory RF center, see Methods). Further stimulus size increases led to an average reduction of responses by 62 ± 3% (mean ± SEM, n = 40) and 77 ± 2% (n = 57) in monkey K and H respectively, i.e. an increase of inhibition or surround suppression (**Figure 1A, S1**), in line with previous observations [26–28]. Importantly, no evidence of rhythmic activity was seen under this condition (**Figure 1B, C**).

**Figure 1:**
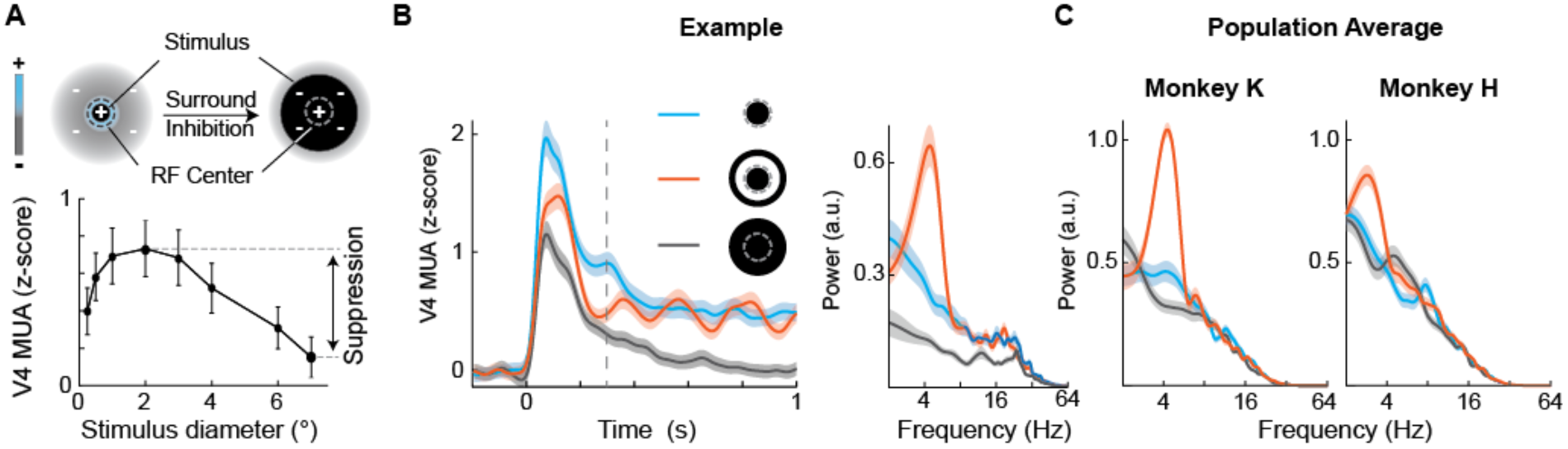
Theta modulation of MUA arises from RF center surround interactions. (A)RF composition with an excitatory center (blue) and a suppressive surround (gray). Small and large stimuli (black disks) are shown to illustrate their relationship with RF structures. Lower panel depicts example size tuning curve from one representative MUA channel from monkey K (suppression index SI = 0.78, based on ∼100 trials per condition). (B) Left panel shows example MUA response from same recording electrode as in (A) to three different stimulus configurations: small 2° disk (blue), large 6° disk (gray), and 2° disk with 4-6° annulus (orange). Vertical dashed line highlights the window for spectral analyses (0.3 – 1s). Right panel depicts corresponding MUA powerspectra (calculated during 0.3-1 s after stimulus onset). Note the presence of rhythmic spiking in the disk-annulus condition only. Shadings around lines depict SEM. (C) Population powerspectra for the same stimuli as in (B), averaged across channels for monkey K (left panel, n = 40) and monkey H (right panel, n = 57). Shadings around lines depict SEM. See also **Figure S1**.

This situation changed profoundly when RF center and surround were stimulated separately using spatially separated visual objects (disk-annulus or disk-flanker stimulation, **Figure 1B, S2**): Only under these stimulus conditions, substantial rhythmic spiking in the theta frequency band (peak frequencies: monkey K: 4.1 ± 0.2 Hz, monkey H 3.4 ± 0.1 Hz) emerged in the MUA time course as well as its spectral representation in most electrode channels of both monkeys (**Figure 1B-C, S1E**): 98%, 39/40 electrodes in monkey K; 79%, 45/57 electrodes in monkey H (see Methods). As shown in **Figure 1C** and **Figure S1E**, across electrodes the strength of theta power increased on average by 185 ± 27.9 % (monkey K) and 158 ± 40.3 % (monkey H) in the disk-annulus condition and therefore significantly more compared to the condition when a single large disk with the same outer diameter was presented (-11 ± 7.8 % for monkey K and 43 ± 23 % for monkey H, *p* = 7x10^-8^, n = 40 and *p* = 2x10^-5^, n = 57, respectively, Wilcoxon signed rank test). Autocorrelograms, another method used to assess rhythmicity, revealed similar results (**Figure S1C**). Examination of the local-field potential (LFP) revealed a very similar pattern in that a prominent theta-peak emerged under disk-annulus stimulation conditions (**Figure S1D**). These results therefore extend previous findings of surround suppression effects in V4 [26,27,30,31,38,39] and demonstrate that placement of multiple objects with respect to neurons’ RF center and surround induces theta-rhythmic activity.

### Theta emerges from RF competition between neighboring neuronal populations

In their recordings from IT neurons, Rollenhagen et al. [14] observed similar theta oscillations as we did and further demonstrated that the initial phase of the oscillation, whether it was inhibitory or excitatory, depended on the order of the stimulus presentation. We confirmed this result at the level of our V4 MUA assessment by dissociating the onset of center and surround stimulation in time (**Figure S2A-C**): Following initial presentation with a stimulus in the RF center, adding flanker stimuli to the surround resulted in a temporary suppression of the response followed by a strong theta rhythmic oscillation. Conversely, when the RF center was stimulated after initial surround stimulation, the theta rhythm started with an initial excitation resulting in an out-of-phase oscillation pattern compared to the previous stimulation condition. Therefore the location of the second stimulus with respect to the first stimulus and the channel’s RF influenced the phase of the resulting oscillation. These phase differences suggest competitive interactions between neighboring neuronal populations. Under our stimulation conditions, one pool of neurons would be excited by the presence of the disk stimulus in their RF, whereas a neighboring pool of neurons would be excited by the presence of one of the flankers in their RF. Disk and flanker representing neuronal populations would inhibit each other (**Figure 2A**). Our recording approach using multi-microelectrode arrays in retinotopically organized V4 allowed us to probe for such RF interactions. To this end we compared the MUA from electrodes with RFs overlapping either the disk or the flanker stimuli, thereby drawing activity from the two stimulus representing neuronal populations (**Figure 2B**). As predicted, the MUA of these electrodes exhibited a simultaneous theta rhythm (**Figure 2B,** see **Figure S2D** for a population powerspectrum), but with a prominent phase offset between electrodes with RF coverage of disk vs. flanker stimulus (**Figure 2C**, monkey K: ΔΦ = 77 ± 1.6° (54 ms at 4 Hz), n = 385, monkey H: ΔΦ = 134° ± 5.7° (93 ms at 4 Hz), n = 20 channel-combinations). In other words, high MUA at one electrode site was accompanied by low MUA at the neighboring site, suggesting mutual inhibition mediated by the RF organization as an underlying mechanism of this rhythm (**Figure 2A**).

**Figure 2:**
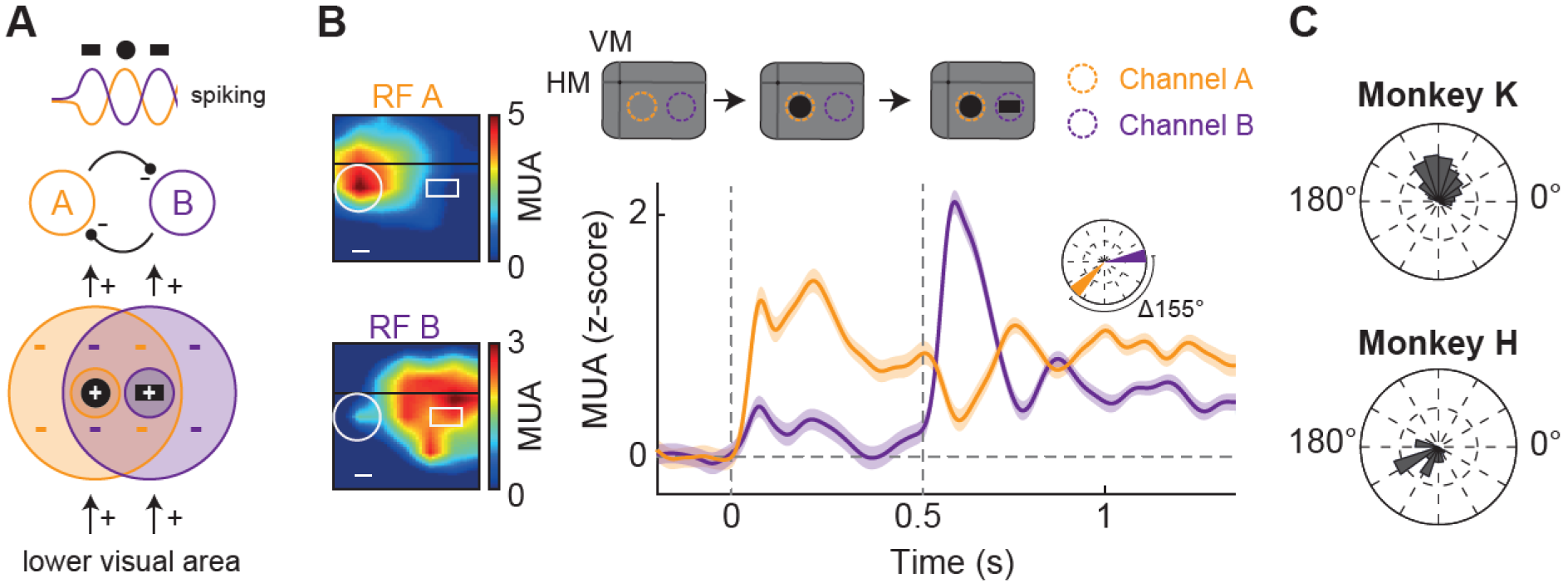
Anti-phasic theta oscillations from nearby electrodes with adjacent RF coverage. (A)Hypothesized mechanism underlying the neural theta oscillation. If two separate stimuli (e.g. disk & bar, simultaneously presented) each drive one population while being in the suppressive surround of the other, the anti-phase oscillation is triggered. (B) Upper panel: orange and purple dashed circles indicate RF of two representative MUA channels. Lower panel shows MUA responses from two example MUA channels from monkey K, each driven either by the disk (orange) or the bar (purple). Inset depicts theta phases of both channels. Left panels show the channels’ RFs with disk and bar stimuli, scale bars signify 1°. Shadings around lines depict SEM. (C) Population distribution of phase-differences between disk‐ and bar-selective channels for both monkeys (for monkey K: mean = 77° ± 1.6°, n = 385 channel combinations; monkey H: mean = 134° ± 5.7°, n = 20 channel combinations). See also **Figure S2**.

### Microsaccades cannot explain the MUA rhythm

What might be the behavioral correlates of this rhythmic spiking? The results presented so far were all obtained in monkeys maintaining passive fixation within 1° radius on a central fixation spot while the stimuli were presented in the periphery. The monkeys were not allowed to carry out saccades to the stimuli and eye movements were continuously monitored. However, this leaves the possibility that small fixational eye movements (microsaccades, MS, **Figure 3, S3**), which could result in brief changes of the visual input to neurons, might have contributed to our effects. MS have been previously associated with attention [40,41] and might at least under some conditions even occur rhythmically [1,19,42] in tight correspondence with neural rhythms [19,20,37]. In our data MS, when they occurred, were quite distinct from the observed theta MUA rhythm: MS were not present in every trial; when MS occurred during a trial after stimulus onset (63% of trials for monkey K, 48% for monkey H), then not at a fixed latency with respect to stimulus onset and/or with an apparent rhythm (**Figure 3A-B).** Across trials we found microsaccades to occur about once per second (1.0 ± 0.1 Hz in monkey K and 1.3 ± 0.2 Hz in monkey H), which is a lower frequency compared to the simultaneously measured MUA rhythm (monkey K: 4.1 ± 0.2 Hz, monkey H 3.4 ± 0.1 Hz) (**Figure 3A-B)**. When they were present, MS triggered a transient MUA response with a peak latency of ∼100-200 ms, consistent with previous findings [20,23,43], that was arrhythmic (**Figure 3C, Figure S3)**. In a final step, we examined MUA during trials during which no MS occurred during stimulus presentation. In the example presented in **Figure 3D**, MS were only present during the baseline period yet with no apparent rhythm in the MUA. No MS were present during visual stimulation, yet the center-surround stimulus clearly evoked theta-rhythmic MUA modulation. The same pattern was present also when examining the average MUA across all trials without MS (**Figure 3E**, upper panel) and when directly comparing theta MUA power in trials with and without MS across electrodes (**Figure 3E**) which showed no significant difference (p = 0.27, n = 40 and p = 0.83, n = 57 for monkey K and H, respectively). MS could therefore not account for the stimulus-induced theta rhythm in our MUA data.

**Figure 3:**
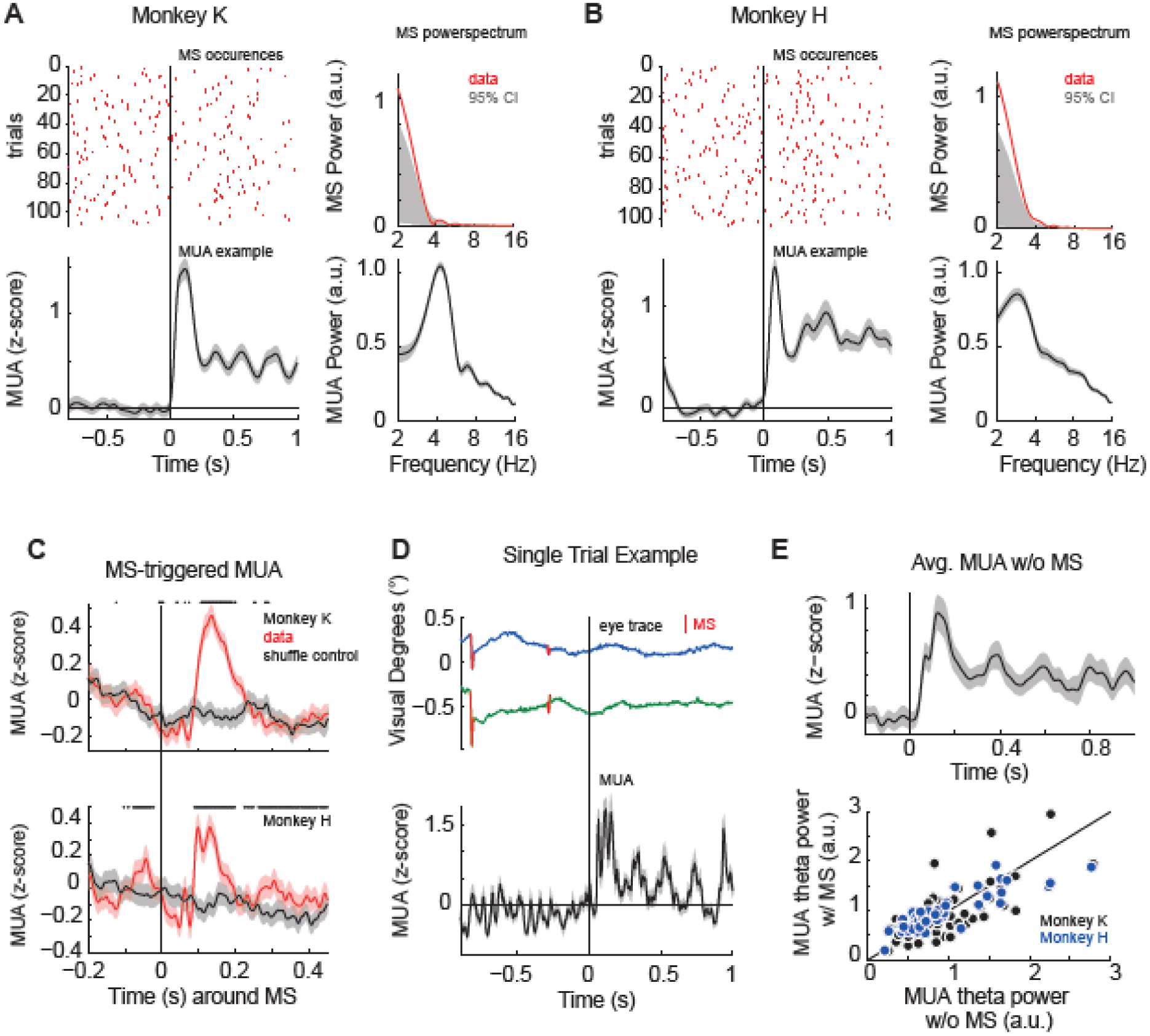
The MUA rhythm does not depend on microsaccades. (A)Raster plot shows microsaccade (MS) occurrences over trials and time, lower panel shows example MUA response (example channel, averaged across trials). Right panels depict powerspectra of microsaccade occurrences and MUA responses (population average across trials and channels, same as in Fig. 1). In 37% (40/108) of the trials no microsaccade occurred in the time period used for the spectral analysis (0.3 - 1s after stimulus onset). (B) Same as (b), but for monkey H. 52% (54/104) of the trials exhibited no microsaccade in the spectral analysis window. (C) MS-triggered MUA from monkey K (upper panel) and H (lower panel). Zero represents the time of MS occurance, the red and gray traces show the actual data and shuffle-control, respectively. Stars on top highlight significant differences between both conditions (*p* < 0.05, Bonferroni-Holm corrected for multiple comparisons). (D)Top panel: single trial example of eye movements of monkey K (x-signal: blue, ysignal: green). Microsaccades are highlighted in red (n = 2). Lower panel shows a single trial MUA response to a disk-annulus stimulus (disk: 2°, annulus: 6° outer, 4° inner diameter) averaged across channels (n = 40). Solid and dashed lines highlight stimulus onset and begin of spectral analysis window, respectively. (E) Upper panel show the average MUA from one example channel from monkey K averaged across trials without MS after stimulus onset. Lower panels compares the theta power per channel based on trials with (‘w/’) and without (‘w/o’) MS for monkey K (black) and H (blue) showing no significant difference (p = 0.27, n = 40 and p = 0.83, n = 57 for monkey K and H, respectively). See also **Figure S3**.

### Rhythmic reaction time fluctuations during distributed attention

Investigations in humans have demonstrated the presence of rhythmic behavior in the theta range (3-9 Hz) in distributed attention tasks involving multiple objects [6,10] suggesting that attention might sequentially sample objects over time rather than statically increase the range of its “spotlight”. We hypothesized that the theta-modulated MUA evoked by center-surround stimulation in the presence of multiple stimuli (passive viewing) might mechanistically underlie behavioral sampling rhythms present under distributed attention. To test this, we trained the monkeys on a task which employed center-surround stimulation in the context of distributed attention. Monkeys had to detect a small luminance change (target) that was randomly displayed on either the RF center (disk) or the surround stimulus (flanker) by executing a saccade to this location (**Figure 4A**, see Methods). Targets were presented after a randomized period within up to 750 ms following stimulus onset, allowing the post-hoc reconstruction of reaction times (RTs) sorted by target onset times across trials. The monkeys could perform this task with ease (monkey K: 92% correct, 7 sessions, 878 correct trials, mean RTs: disk: 212 ± 6.6 ms, flanker: 220 ± 3.8ms; monkey H: 96% correct, 8 sessions, 1509 correct trials, mean RTs: disk: 210 ± 6.9 ms, flanker: 180 ± 6.0 ms). By analyzing the reaction times as a function of target onset times, we found that their behavior was not constant across target delays, but rather fluctuated over the assessed time period. For example monkey K, at the time point highlighted by the first grey bar in **Figure 4B** responded faster to the target when it occurred on the center disk stimulus than when it occurred on the nearby flanker stimulus. Thus it seems that the monkey’s attention was likely focused on the disk stimulus at this time requiring re-orienting when the target instead occurred on the flanker stimulus. Importantly this bias towards the disk stimulus was periodic and alternated with periods favoring the competing flanker stimulus resulting in the overall RT fluctuation over time. We assessed the rhythmicity of these fluctuations by computing their powerspectrum and found a significant peaks at 4.3 Hz in monkey K and 5.7 Hz in monkey H for the center stimulus (theta frequency range, **Figure 4B** upper right panel**, S4A**, red lines, *p* = 2x10^-4^ and *p* = 2x10^-4^ for monkey K and *p* = 2x10^-4^ and *p* = 0.01 for monkey H for the center (red line) and flanker target (blue line), respectively, randomization test, n = 5000). This assessment also confirmed that periods of short RTs for one target location coincided with RT costs for the competing target location (theta phase-difference: 95° and 97°, equaling a 66 and 67 ms shift at 4 Hz, in monkey K and H, respectively). Thus monkeys, similar to humans [4,6,10], appear to engage in rhythmic attentional sampling under conditions when multiple stimuli compete for perceptual selection.

**Figure 4:**
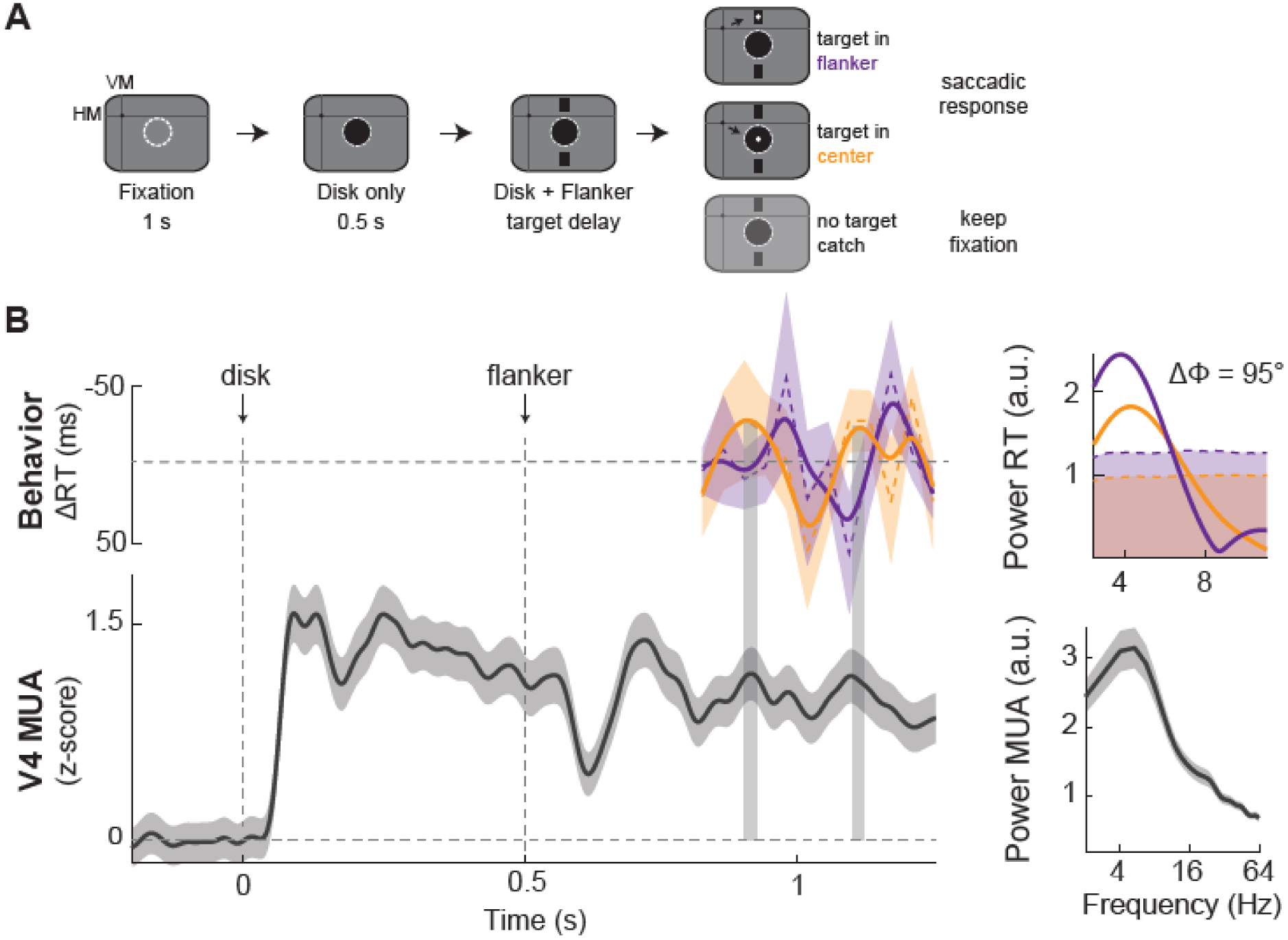
Same theta (3-6 Hz) rhythm in MUA and reaction times during distributed attention task. (A)Experimental design. VM/HM = vertical/horizontal meridian. The luminance change that had to be detected by making a saccade to its location could appear either in the in the center (orange) or flanker (purple, distributed attention task). 1/3 of the trials constituted of catch trials where no target appeared. (B) Upper trace shows the reaction time (RT, normalized as deviation from mean) from monkey K for distributed attention conditions. RT to the center target position is shown in red, to the flanker target position in blue. Non-smoothed RT data shown as thin dashed lines. Lower trace shows MUA response recorded during catch trials (no target appearing; during distributed attention). Vertical gray shaded areas highlight the peak-to-peak locking between the behavioral and neural oscillation (RT to center location (orange) and MUA with corresponding RF). Right panels depict powerspectra of the respective rhythms and the RT phase difference between center and flanker conditions. Shaded areas of RT powerspectra highlight 95% confidence intervals based on shuffled surrogate data. Shadings around other lines depict SEM. See also **Figure S4**.

### Attentional sampling is coupled to rhythmic V4 MUA

Analysis of the neuronal data accompanying the behavior during the distributed attention condition revealed the following pattern: After initial excitation through the disk in the RF center, addition of the flankers initiated a suppression in spike rate that was reliably followed by a rhythmic oscillation pattern at theta frequencies (**Figure 4B** lower panel, also **Figure 6A, S5A**), similar to the rhythmic fluctuations in RTs. Moreover, when aligning the two signals in time, their relationship became clear (**Figure 4B**, lower panel): Periods of increased MUA during the oscillation were temporally aligned with periods of shorter RTs, whereas lower MUA levels were associated with longer RTs. To quantify this relationship between the RT and neural oscillations, we computed the cross-correlation between the RT time course across trials to the center target and the MUA recorded during catch-trials (**Figure 5A**). We found significant correlations in 36 of 41 (88%) and 40 of 57 (70%) electrodes (*p* < 0.05, randomization-test, see Methods) with mean correlation coefficients of 0.41 ± 0.01 (n = 36) and 0.42 ± 0.01 (n = 41) for monkey K and H, respectively (Figure 5A). Cross-correlations were also prominently rhythmic in the theta range (peak frequencies: 4.6 ± 0.18 Hz, n = 36 and 5.9 ± 0.29, n = 41 for monkey K and H, respectively), indicating rhythmic synchronization between behavior and neural activity (**Figure 5B,** significantly theta-rhythmic RT-MUA combinations: 36/36 (100%, monkey K) and 40/40 (100%, monkey H), *p* < 0.05, randomization test).

**Figure 5:**
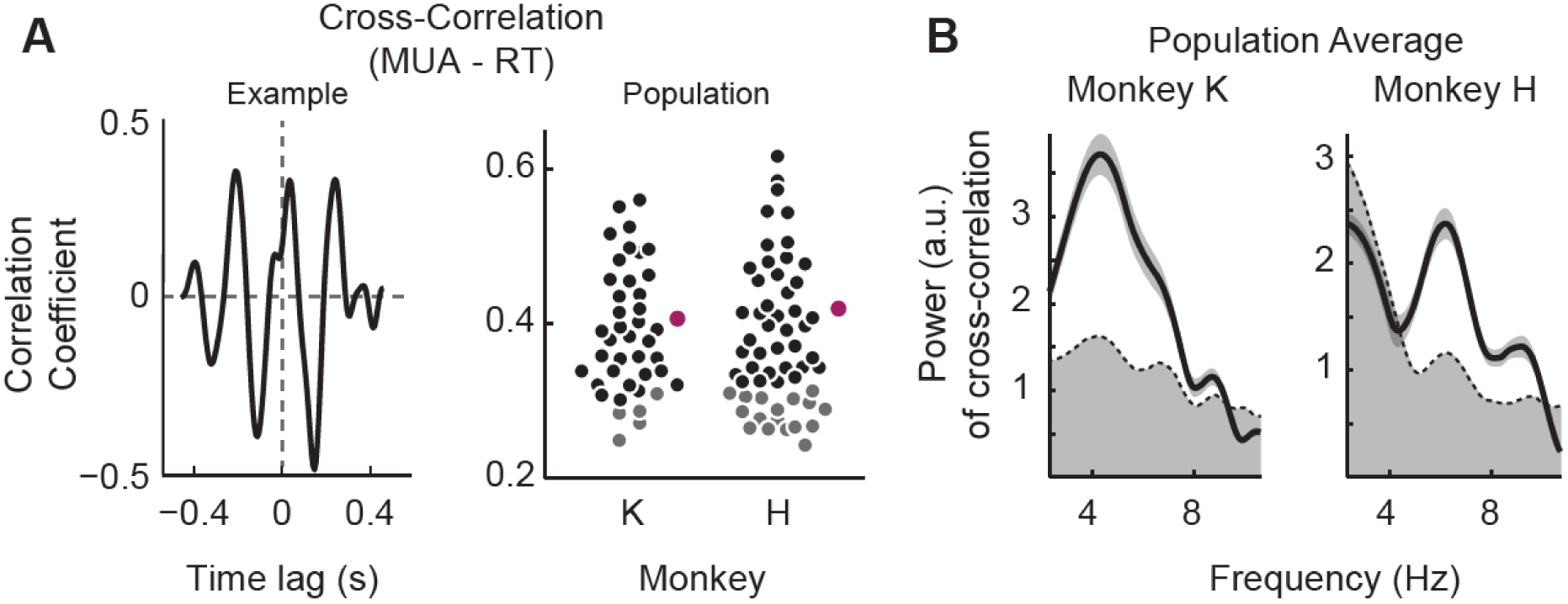
Rhythmic distributed attentional sampling is correlated with cortical MUA rhythm. (A)Example cross-correlogram between RT and MUA from monkey K during distributed attention (left panel). Right panel shows the distributions of correlation coefficients of RT-MUA channel combinations for monkeys K (left, n = 41) and H (right, n = 57). Electrodes with significant cross-correlations to RT are shown in black (non-significant channels in gray). The mean MUA-RT cross-correlations across significant electrodes are shown in purple for each monkey. (B) Powerspectra of cross-correlations for monkey K (left) and H (right) averaged across channels. Shaded area below dashed line depicts 95% confidence interval. Note the significant cross-correlation at theta frequencies.

**Figure 6:**
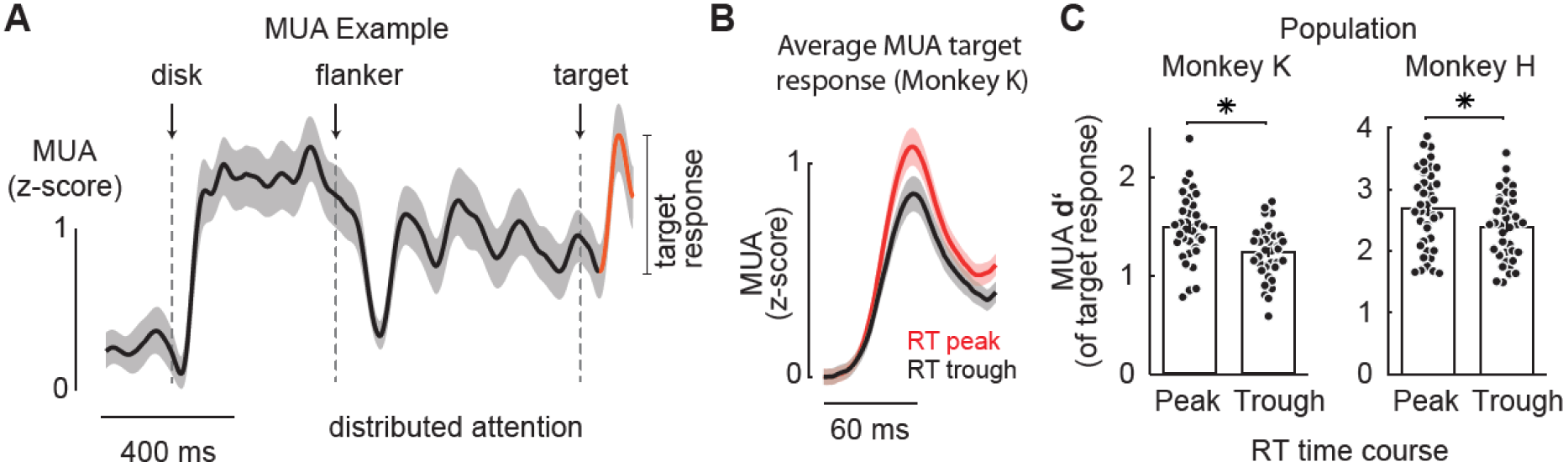
Strength of MUA response to target in relation to periodic fast and slow RT. (A)MUA from one representative electrode (monkey K) during trials in which the target appeared over the disk stimulus (center position). The MUA response to the target’s luminance change is highlighted in orange on the background of the stimulus induced oscillation (black). (B) Average MUA target response across channels comparing response strength during target delays that corresponded to peaks (red) and troughs (black) of the RT time course. (C) Population distributions of d’ of MUA target responses across channels for RT peaks and troughs for monkey K (left) and H (right) during distributed attention. Asterisks denote significant differences. Shading and errorbars depict SEM. See also **Figure S5**.

### Faster reaction times are preceded by larger responses to target stimulus

So far the analysis focused on the rhythmic relationship between RT time courses and MUA collected during interspersed catch trials. This was enabled by consistently phase-locked MUA time-courses induced by the center-surround stimulus configuration. This analysis revealed the intimate relationship between the behavioral and MUA rhythms across trials. In the next step, we chose a different strategy and instead aimed to directly assess the trial-by-trial representation of the target, i.e. the luminance change, during the course of the observed theta rhythm in the RT. Our rationale was that the representation of the target would be affected by the state, i.e. the phase of the underlying MUA rhythm, and therefore translate into corresponding RTs. More specifically we reasoned that during the peak periods of the rhythm, neurons might be more sensitive to the target, therefore respond with greater amplitudes and result in shorter reaction times. In other words, if the behavioral modulations were conveyed by rhythmic changes of neuronal excitability, the response of neurons to the physically same target stimulus should differ in strength between fast and slow RTs. To directly test this for the data collected during the distributed attention task we quantified the MUA responses to the target stimulus during peak vs trough periods of the RT time course (**Figure 6A**). **Figure 6B** shows the average MUA responses to the target across channels (n = 35) recorded during RT peaks and troughs from monkey K. To further quantify this modulation across channels, we normalized these MUA target responses to a MUA baseline that preceded the response by 60 ms, resulting in d-prime (*d*’) as a measure for the neuronal sensitivity to represent the target (see **Figure S5B** for a sketch illustrating the calculation of *d*’). Indeed, we found that *d*’ during peaks of the behavioral oscillation (fast RT period, see Methods for a detailed description) was significantly larger than that during an oscillation trough (**Figure 6C**, *p* = 2.7x10^-6^, n = 35 in monkey K; *p* = 4.4x10^-6^, n = 41 in monkey H, see Methods). Therefore, as predicted, trials with stronger responses to the target stimulus resulted in shorter RTs. Due to the tight link between behavioral and MUA rhythms this indicates that theta rhythmic activation influenced the neuron’s sensitivity to respond to the target and therefore mediated the RT rhythm.

## Discussion

We found that receptive field center surround interactions of V4 neurons induce prominent MUA at theta (∼3-6 Hz) frequencies in the presence of multiple visual objects. This theta rhythm was stimulus-induced and occurred irrespective of the presence or absence of microsaccades. When monkeys had to simultaneously monitor the visual objects (distributed attention) to detect an unpredictable luminance change (i.e. the target) in one of the objects, reaction times were modulated by a similar theta rhythm as in the MUA. Correlation analyses confirmed a significant locking of RT fluctuations to V4 MUA. Furthermore, the strength of the MUA response to the target change within the theta rhythm was predictive of the monkeys’ periodic fast and slow reaction times. In the following we discuss these findings with respect to generative mechanisms of theta oscillations in visual cortex and their function in the context of attentional sampling.

### Theta rhythm in visual cortex: effects of stimulus competition and microsaccades

Assessment of single‐ or multiunit activity from visually responsive parts of the brain has resulted in select number of studies that report theta-rhythmic spiking. Spiking in the theta range has been observed in the pulvinar [44] and mediodorsal thalamus nuclei [45,46], and at the cortical level in V4 [13] and inferotemporal (IT) areas [14–18,47] of awake non-human primates. In addition theta rhythms were reported in a local field potential recordings assessing the role of oscillations for inter-areal communication [19,48–51]. What are the mechanisms that contribute to the emergence of theta rhythmic activation? In what follows we discuss how microsaccades and stimulus competition contributed to the neuronal rhythm in our data and how our findings tie-in with the existing literature.

Microsaccades (MS) are often observed in the context of attentive, explorative behavior. Recent evidence indicates that at least under some tested conditions, they may occur rhythmically and link to rhythmic neuronal activation [e.g. 11,14]. In both studies MS occurred with a higher likelihood every 250-300 ms, i.e. roughly corresponding to a 3-4 Hz theta rhythm. These theta-rhythmic MS occurrences modulated the power of the faster gamma band in the LFP. We therefore tested whether MS during our paradigm exhibited a similar theta rhythm that could account for the theta rhythmic neuronal modulation in our MUA data. Under our stimulation conditions MS occurred at a lower rate than in the Lowet and Bosman studies, namely around 1 Hz, consistent with many other reports in the literature [1,52]. MS rates are influenced by a number of stimulus-related and cognitive factors that likely explain the variation across studies [52]. For example, microsaccade rate is known to increase as a function of fixation duration [1]. In the study of Lowet et al. the time period that was used for the microsaccade analysis was after at least 1.8 seconds of visual stimulation. Similarly, in the Bosman et al study monkeys maintained fixation for several seconds. In our paradigm under passive fixation the stimulus duration was 1 second and therefore considerably shorter, which likely contributed to the lower MS rate in our data. The 1 Hz MS rate in our data was lower than the 3-6 Hz rhythm in the MUA and therefore seems to reflect a different process. MS resulted in a MUA response with peak latency between 100 and 200 ms, consistent with previous findings in V4 [20,23]. Whereas MS occurred not in every trial and at different times within a trial, theta-rhythmic MUA occurred highly consistent across trials, even in the absence of MS. While our results and similarly the study by Rollenhagen et al. based on recordings in IT did not show any relationship between the MUA rhythm and MS occurrence during short stimulus presentations, it is conceivable in particular for longer stimulus periods that the stimulus-induced MUA rhythm could trigger the occurrence of rhythmic microsaccades and overt sampling or that microsaccades could reset the phase of an ongoing theta rhythm.

In our study theta-rhythmic MUA was triggered by the presence of multiple stimuli with respect to the neurons’ RFs. This result provides a mechanistic explanation for previous findings with theta rhythmic activity in visual cortices [13–18] on the basis of center-surround RF interactions (**Figures 1, 2**). Increasing the stimulus beyond the RF or adding a second stimulus to the one presented in-side the RF both decreased the initial stimulus response as expected from previous results [30] and a sensory normalization mechanism [53,54]. In line with a reduced suppression drive, the initial stimulus response was intermediate when instead of a single large stimulus, only an annulus or bar was shown outside the RF. It is under these intermediate excitation conditions in the presence of two stimuli, that MUA became rhythmic (**Figure 1-2**). The oscillation signal was initially negative when the surround (flanker) stimulus was added to the central (RF) stimulus (due to the incoming suppression) and positive under the opposite order, consistent with previous findings [14]. In our V4 array recordings we could also observe this effect for nearby electrodes likely sampling from neighboring neuron pools (**Figure 2**) providing evidence for competition induced theta oscillations in visual cortex occurring at intermediate excitation levels.

Modeling work by Moldakarimov et al. suggested that the emergence of slow oscillatory activity is influenced by the strength of inhibition between contributing neuronal pools and the timing of spike fatigue and synaptic delay time [35]. Previous experimental research demonstrated that brief (50 ms) presentation of a second stimulus outside the RF in addition to a stimulus inside the RF induces surround suppression in V4 neurons 75 – 235 ms following stimulus onset, i.e. in the range of a slow theta cycle [30]. Furthermore, increasing the stimulus contrast or directing attention to the surround stimulus was effective in increasing the suppressive effect on the neurons’ responses. It appears that our stimulation conditions prolonged this surround suppression mechanism resulting in theta-rhythmic activation. Variability with respect to RF coverage of the stimulus as a function of eccentricity, and consequential latency differences of horizontal connections, is likely the source for observed frequency and phase variations in our data. Future work will need to clarify how the local connectivity contributes to surround suppression and theta-rhythmic spiking, including whether the mechanism is truly inhibitory or better accounted for by a release of excitation. Furthermore there is a need to understand how other factors such as the number of stimuli and distance between them affects theta. Finally it will be important to determine whether similar receptive-field competition mechanisms are also underlying the emergence of theta oscillations in other cortical areas with different RF coverage and in the context of other selection tasks such as during binocular rivalry [16] and working memory [13,49].

### Theta rhythms during spatial attention

Theta rhythmic fluctuations of perceptual performance are described in a rapidly growing literature. In the presence of multiple stimuli the “attentional focus” appears to switch between potentially relevant objects resulting in out-of-phase behavioral measures of attention (e.g. performance, reaction times) towards these stimuli [6,9–11,55]. The present data extend the evidence that such attentional sampling exists in humans to macaque monkeys. Comparable behavioral results in monkeys were presented by the Kastner group [56].Both species appear therefore to engage in theta-rhythmic sampling when subjects are asked to covertly track two or more objects and respond to a spatially and temporally uncertain change in one of the objects. An important aspect to temporally “capture” the locus of attention across trials and subjects is to reset ongoing performance fluctuations to an external event [57,59]. For example, Landau et al reset attentional performance to the onset of a mask stimulus surrounding one of their targets. In our paradigm the sequential presentation of the two stimuli induced a reset both at the behavioral as well as at the neuronal level. This enabled us to track subsequent rhythmic fluctuations and cross-correlate their time courses in MUA and reaction times.

While neural rhythms at the EEG/MEG/LFP level have been described in the presence of a wide variety of tasks, the specific relationship between behavioral and neural rhythms is less clear. Several studies demonstrate that neural rhythms before target onset determine performance (e.g. [4,7,58]). Recently Jia et al demonstrated that attention-related EEG alpha oscillations were modulated every 200 ms during distributed attention and that alpha correlated with performance on the unattended object [60]. Furthermore it has been shown that gamma oscillations in response to their stimulus were modulated by theta and that the phase of this theta-gamma coupling was predictive of detection accuracy [11]. Our results show that a similar theta rhythm is present both at the neural level in MUAs and LFPs as well as in behavioral performance. The presence of two stimuli was associated with a neural theta rhythm where the phase of this neural theta predicted location specific performance, such as theta “up-states” where associated with shorter reaction times and theta “down-states” with longer reaction times. We further showed that these rhythmic states modulated the amplitude of the target-evoked response, so that larger evoked responses resulted in faster reaction times. This is consistent with a wide set of findings that the phase of oscillations is predictive of performance and reaction time fluctuations for near-threshold stimuli ([8,58,61–64], however see [65,66]) and reflect fluctuations in neural excitability [59,67,68].

How distributed attention affects the responses of single neurons in visual cortex is not well understood. Addressing this at the level of V4 a recent study found reduced responses during uncued (distributed) vs cued (focused) attention conditions [69]. By recording from both hemispheres simultaneously the authors examined whether they could find any evidence for a switching of responses that may account for attentional sampling of the target locations from both hemifields. The authors did however not observe any signs of a switching mechanism in their data. Methodological aspects, such as a short 200 ms analysis window and the absence of a reset event to capture attention may explain the difference to our results. Moreover it is possible that no rhythmic activity was observed as stimuli were spread across hemifields. It may therefore be that local inhibitory input from the same hemifield is necessary to elicit theta rhythmic activation in V4.

Our neurophysiological results of theta rhythmic MUA in V4 provide a direct correlate for theta-rhythmic attentional sampling. We demonstrate that center-surround interactions are at the origin of this theta rhythm in V4. Excitation of one population during the theta cycle facilitates transmission of stimulus information from its RF, while at the same time information from the RF location of the neighboring population is suppressed. The duration of a theta cycle (∼200 ms) would allow for sufficient time for perceptual processing of at least one object [70] in addition to generating a behavioral reaction, such as a saccade to it. The succession of theta cycles as part of the sampling rhythm may thus act as a dynamic selection mechanism in visual cortex to ensure efficient perceptual processing, but may extend also to other parts of the brain in the context of spatial navigation and memory [71–74], and therefore constitute a fundamental aspect of brain function.

## Acknowledgments

This work was supported by Emmy Noether grant 2806/1 and ERC starting grant “OptoVision” to M.C.S. We thank all ESI staff, the directors Drs. Pascal Fries and Wolf Singer, in particular Hanka Klon-Lipok, Sabrina Wallrath, Michael Stephan, Kai Roennburg, and Johannes Boldt and Axel Klug of MPI Tübingen for their technical support. We are grateful to Drs. Alexander Maier, David A. Leopold and Alex Thiele for very helpful discussions and comments on a previous manuscript version. The authors declare no conflict of interests.

## Author contributions

Conceptualization, R.K., J.T.S., M.C.S.; Methodology, R.K., J.T.S., K.A.S., M.C.S..; Software, R.K., J.T.S., K.A.S., K.K.; Formal Analysis, R.K..; Investigation, R.K., K.A.S., K.K.; Writing - Original Draft, R.K. and M.C.S.; Writing - Review & Editing, R.K., J.T.S., K.A.S., M.C.S.; Supervision, M.C.S..; Funding Acquisition, M.C.S.

## Declarations of interest

The authors declare no competing interests.

## Methods

### Subjects

Two healthy adult male rhesus monkeys (*Macaca mulatta*, monkey K and H) participated in the study. All procedures were approved by the Regierungspräsidium Darmstadt and carried out in accordance with the applicable laws and regulations according to EU directive 2010/63. The monkeys were group housed in enriched environments of the animal facility and with access to outdoor space. All surgeries were carried out aseptically under gas anesthesia using standard techniques including peri-surgical analgesia and monitoring. Each monkey received a titanium-made head-immobilization implant and a recording chamber in addition to Blackrock multi-electrode arrays including a connector implant (Blackrock Microsystems, Hannover, Germany). Throughout the study animal welfare was monitored by veterinarians, technicians and scientists.

### Behavioral task

During all the experiments eye movements were tracked using an infrared eye tracking system at a sampling rate of 500 Hz (EyeLink 1000, SR research, Ottawa, ON, Canada). All stimuli were presented on a Samsung 2233RZ LCD screens (120 Hz refresh rate, 1680 x 1050 resolution, viewing distance was 77 cm for monkey K and 86 cm for monkey H). Stimulus presentation and monkey behavior during the experiments were controlled and monitored using MonkeyLogic [75].

For the passive viewing task monkeys were required to keep fixation on a small (0.07° radius) white dot during the entire trial. Usually 1 s of fixation baseline was followed by 1.5 s of stimulus presentation. When two stimuli were presented sequentially, the presentation length of the first stimulus was 0.5 s, while the second was on for 1 s. The intertrial interval was 1 s for all tasks.

For the attention task, monkeys had to fixate a small central white dot within 1° radius (fixation window) during the presentation of the stimuli. After 1 s of fixation baseline a black disk (2° diameter) appeared in the V4 RF. 500 ms later two flanking, vertically orientated bars, each 1° in length and 0.25° in width, appeared on the screen (1° gap between disk and each bar). After flanker onset, a small target (0.2° diameter luminance increase) was flashed on either the central disk or the upper bar (distributed attention) in varying delays. Timing of the target relative to the flanker onset was randomized across trials within a 750 ms time window starting directly after flanker onset (20 x 37.5 ms time points, linearly spaced across the 750 ms resulting in ∼26 Hz sampling resolution). In order to receive a liquid juice reward the monkeys had to report their detection of the target by executing a single saccade to the location where the target was flashed within a 1.5° radius window. Target contrasts were chosen so that the monkey’s mean performance was >= 90% (contrast as luminance ratio of the target relative to the black stimulus: 0.04 and 0.15 (monkey K), 0.07 and 0.18 (monkey H) for the center and flanker target, respectively). Due to a masking effect arising from the flanker onset and consequential low performance we excluded the first 250 ms of the target delay spectrum from further analysis, resulting in a 500 ms RT analysis window.

In addition, catch-trials were randomly included during a third of all trials. Their purpose was to record the stimulus induced oscillations uninterrupted by saccadic responses and to monitor and test monkeys’ behavior. During these trials flanker stimuli were presented in addition to the disk stimulus for at least the duration of the longest possible trial in detect conditions without the occurrence of any luminance changes. This ensured that catch trials could not be identified by the monkeys until the end of the trial. In order to receive the reward monkeys had to keep fixation during the entire catch-trial. Otherwise, the timing of stimulus events and the stimulus parameters were identical to the other trials.

### Spectral Analysis

To obtain spectral information of multi-unit activity and reaction times, we performed a spectral analysis using a Hanning-tapered Fourier transformation. Visual inspection of the spectra revealed a prominent peak in the 3-9 Hz which is referred to as theta throughout the paper. The phase of the oscillation was extracted from the complex Fourier coefficients using the same parameters as above. Spectral peaks were identified as band-limited peaks in powerspectra as judged by visual inspection for each monkey independently.

Average values of power within frequency windows were calculated as the mean value across time and frequency. Differences between conditions were tested with nonparametric Wilcoxon signed rank tests (paired data), Mann-Whitney U tests (unpaired data) or computational randomization statistics. Values in the text are always mean ± SEM unless stated otherwise.

To test for significant oscillations we took the following approach: Since we were interested in neural dynamics that are defined by their specific time-dependent modulations we computed surrogate data that specifically destroyed the timing information of the measured experimental data. The surrogate data was generated by randomly time shuffling the reaction time courses (detrended, non-smoothed), i.e. randomly re-assigning a given value of the RT vector a new position (index) in the vector. This was repeated 5000 times. In a final step, the very same analysis as for the actual data (spectral analysis, cross-correlation) was performed on the surrogate data (i.e. 5000 time-shuffled trials). From these randomization procedures the p-value corresponds to the proportion of times for which the power in the surrogate data exceeded the power in the actual experimental data.

For the quantification of reaction time course oscillations surrogate data were computed on the non-interpolated RT time courses.

### Analysis of behavioral data

Behavioral data were processed and analyzed using custom-written code for MATLAB (MathWorks, Inc.) and the MATLAB toolbox FieldTrip [76]. Only correct trials were used for the analysis. Reaction time was measured as the time between target presentation and the eye position leaving the fixation radius (1° radius). For the experiments studying the center-surround interactions under passive fixation (**Figure 1, S1**) 4 sessions (3358 correct trials in total) in monkey K and 4 sessions (4146 correct trials in total) in monkey H were recorded, i.e. ∼105 correct trials per condition (as used in **Figure 1, S1**).

For the paradigm with a sequential presentation of disk and bar under passive fixation (**Figure 2**), 3 sessions (60 correct trials per condition) in monkey K and 2 sessions (30 correct trials per condition) in monkey H were recorded. For the disk flanker timing experiments under passive fixation (**Figure S2A-C**) 2 sessions (41 correct trials per condition) for monkey K and 1 session (20 correct trials per condition) for monkey H were recorded.

With the detection paradigm, we recorded 7 sessions in monkey K (878 correct trials in total, 92% performance, mean RTs: disk: 212 ± 6.6 ms, flanker: 220 ± 3.8ms) and 8 sessions in monkey H (1509 correct trials in total, 96% performance, mean RT disk: 210 ± 6.9 ms, flanker: 180 ± 6.0 ms).

Due to the relatively high proportion of catch-trials and timing conditions, reaction time data were pooled across sessions after being normalized session-wise as deviation from mean (resulting in ∼20 and ∼30 correct trials per target delay for monkey K and H, respectively).

To obtain RT timecourses as a function of the variable target onset times across trials (**Figure 4, S4**), we calculated the mean and standard error of the mean RT for each target onset time across trials. This results in a vector of mean RTs with an entry for each target onset time. This vector was then detrended using a second order polynomial and smoothed (for display purposes and the computation of RT powerspectra) using a smoothing spline. Power and phase of the RT timecourse was then calculated using a Hanning-taped Fourier transformation.

### Microsaccade analysis

During visual stimulation monkeys were not allowed to perform saccadic eye movements (except for target detection). Nevertheless, during fixation small eye movements (<1°) can occur. To detect those, we used an algorithm similar to the one introduced by Engbert and Kliegl [77], where microsaccades are detected as outliers in a velocity-space. Detection threshold were set to 5 standard deviations of the velocity distribution. In order to perform a spectral analysis of microsaccade occurrences microsaccade-time points detected in the same time window used for the MUA spectral analysis (0.3 – 1s after stimulus onset, highlighted in **Figure 3** by dashed line) were pooled across trials and used to compute a probability density estimate based on a normal (kernel) function (bandwidth = 0.18). This was then used to compute the powerspectra in the same way as for the MUA.

To compute MS-triggered MUA the time-stamps of MS were extracted and then used a trigger to align MUA across trials per channel. The shuffle control data (Figure 3C, gray lines) was computed by performing the same analysis but using MS timings and MUA from separate trials. The power of MS-triggered MUA was calculated in the time-period following the MS. P-values in Figure 3C were computed across trials using the trial-averaged MUA and corrected for multiple comparisons using the Bonferroni-Holm correction.

### Neurophysiology and Data Preprocessing

Neurophysiological data was recorded from 64 channel Blackrock multi-microelectrode “Utah” arrays (Blackrock Microsystems, Hannover, Germany) that were chronically implanted in the left hemisphere’s area V4 (prelunate gyrus). Electrodes had lengths of either 0.6 or 1 mm arranged in alternating sets of two rows of short and long electrodes. Each electrode was 400 μm away from its neighboring electrodes. Reference wires were inserted over parietal cortex and cerebellum. Neural data was recorded at a sampling rate of 30 kHz using a Blackrock Microsystems Cerebus system.

All neurophysiological data were processed and analyzed using custom-written code for MATLAB (MathWorks, Inc.) and the MATLAB toolbox FieldTrip [76]. The continuous recordings were separated into trials using digital event markers and aligned on stimulus onset. We focused our analyses on the sustained periods excluding the transient onset response (300 - 1000 ms after stimulus onset).

An estimate for multi-unit activity (MUA) was obtained from the high frequency envelope: MUA was extracted from the raw data by high-pass filtering (300 Hz), rectification, low-pass filtering (120 Hz) and then downsampling to 500 Hz [36,78,79]. Data from microelectrode arrays were z-scored per session and then pooled across sessions considering the data as dependent across sessions.

In order to assess the stimulus-specific effects of neuronal spiking multi-unit activity (MUA) data were z-scored channel-wise based on the average prestimulus fixation period (700 ms before stimulus onset). MUA is therefore expressed as z-score values throughout the paper. Receptive fields were mapped for each electrode channel by quantifying their responses to small gratings with a Gaussian mask (1.5° diameter, 100% contrast, moving upward or downward with 1.5 cycles/s, spatial frequency 1.5 cycles/°) presented at 63 positions for monkey K (34 for monkey H) in the lower right quadrant in the range x=1 to 5.5° and y=-4 to 2° for monkey K (x=0.5 to 3.5° and y=-5 to -0.5° for monkey H). The resulting response matrix was convolved with a gaussian window. The receptive field focus was defined as the position eliciting the maximal response.

To be included into further analyses the Euclidian distance between a channel’s RF focus and the stimulus center had to be less than 1.5° (41 and 57 channels in monkey K and H, respectively). This ensured that stimuli were presented into channels’ excitatory center. Mean RF centers were 2.9 ± 0.1 (x-axis) and -0.8 ± 0.1 (y-axis) and 1.4 ± 0.1 (x-axis) and -1.8 ± 0.1 (y-axis) in monkey K and H, respectively.

### Surround Suppression in MUA

We defined the RF center as the diameter of the stimulus that elicited the strongest average activation (across trials and post-stimulus time) in a given channel. To quantify the maximal suppression induced by stimulating the RF inhibitory surround, we identified the stimulus with a larger diameter than the RF center that elicited the minimal average activation within the tested stimulus sizes and computed their relationship as follows:

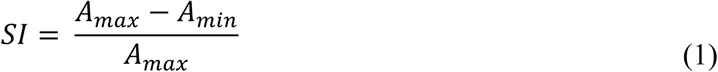

where SI is the Suppression Index, *A*_*max*_ is the maximal and *A*_*min*_ the minimal MUA. The maximal response had to be significantly larger than baseline activity (40/41 channels in monkey K, 57/57 channels in monkey H). By visual inspection, 15% (n = 6/40) and 33% (n = 19/57) channels showed asymptotic suppression in monkey K and H, respectively (**Figure S1)**. The mean size of the center summation RF across channels was 1.7° ± 0.2° (n = 40) and 0.9° ± 0.1° (n = 57) in monkey K and H, respectively.

To quantify the change in theta power as a function of suppression increase (**Figure S1)**, we computed a theta index either between theta power in response to a 2° disk vs. larger disks (3°,4°,6°,7°) or of larger disks (3°,4°, 6° 7°) and disk-annulus stimuli with a 2° central disk and varying annulus sizes (6°,3°; 6°,4°; 7°,3°; 7°,4°; 7°,6°; outer and inner annulus diameter respectively). For the latter comparison the outer diameter of the annulus always matched the large disks in size to ensure that both stimuli reached into the surround by the same distance. The theta index was computed as follows:

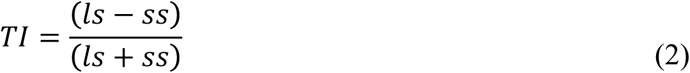

where *ls* and *ss* are the theta powers in response to a large and small stimulus, respectively. In case of the disk vs. disk-annulus comparison, *ls* and *ss* are the theta powers in response to a disk-annulus and corresponding large disk stimulus, respectively.

### MUA phase analysis between electrodes

For the phase analysis displayed in **Figure 2B-C** electrode channels were selected and grouped depending on whether MUA response was stronger for the disk or the flanker stimulus position. In addition, theta power in the post-stimulus period excluding the transient had to be significantly larger than that during the fixation baseline and contain significant theta oscillatory power. This ensured that we performed the phase analysis only on channels that showed significant theta oscillations and were (relatively) selective to one of the two stimulus positions. Note that this selection of channels with significant theta oscillations was only performed in this particular analysis. In all other analyses channels were selected independently of theta power.

### Cross-Correlation between RT and MUA

In order to quantify the relationship between the theta modulation in behavior and MUA, we computed cross-correlations between the mean reaction time time-trace and the MUA signal of each channel for both monkeys as follows [80]:

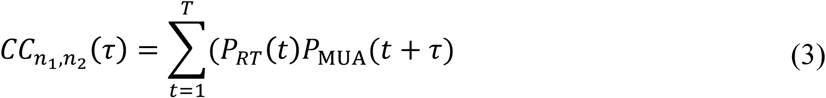

where *T* is the number of discrete time bins, *t* is time, *P*_*RT*_ and *P*_*MUA*_ are the normalized reaction times and MUA PSTHs (averaged across trials, mean subtracted), τ is time lag and *CC*_*n1,n2*_ is the cross-correlation. Before computing the cross-correlation we inverted the reaction times (as seen in **Figure 4B**) such that a positive cross-correlation signifies a coincidence of a high MUA response and fast reaction times.

Before performing the cross-correlation both the RT (detrended, but non-smoothed) and MUA signals were resampled to 150 Hz to ensure that similar time points of both signals are used for the computation. We used time lags between –450 ms and +450 ms. Crosscorrelograms were then normalized such that the correlation coefficient of the autocorrelation at zero lag equals 1.

Correlation coefficients were extracted from the cross-correlations by identifying the maximum value within ±125 ms around 0. To test for a possible rhythmic modulation we computed powerspectra of the cross-correlations (see “Spectral Analysis” for further information) based on the actual data and on those computed from the surrogate data (see above).

### Strength of MUA response to target for fast vs slow RT periods

To quantify the strength of the MUA response to the target (luminance change) with respect to baseline we computed the d’ as a measure of sensitivity as follows:

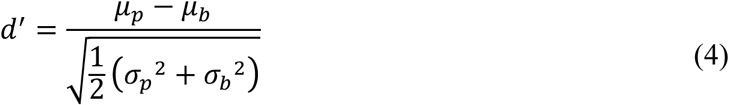

where ***μ*_*p*_** and ***μ*_*b*_** are the means and ***σ*_*p*_,*σ*_*b*_** the standard deviations for the MUA of the peak and the baseline preceding the peak by 60ms, respectively. This analysis was performed on MUA recorded during detection trials (distributed attention condition) during target delay conditions associated with a peak in the reaction time course (i.e. fast RT; 0.9 and 1.1 s for monkey K and 0.85 and 1.02 s for monkey H) vs. target delay conditions associated with a trough (slow RT; 0.82 and 1.01 s for monkey K and 0.78 and 0.98 s for monkey H). Peaks were detected for each condition and channel separately.

